# Drug repurposing screens reveal FDA approved drugs active against SARS-Cov-2

**DOI:** 10.1101/2020.06.19.161042

**Authors:** Mark Dittmar, Jae Seung Lee, Kanupriya Whig, Elisha Segrist, Minghua Li, Kellie Jurado, Kirandeep Samby, Holly Ramage, David Schultz, Sara Cherry

## Abstract

There are an urgent need for antivirals to treat the newly emerged SARS-CoV-2. To identify new candidates we screened a repurposing library of ~3,000 drugs. Screening in Vero cells found few antivirals, while screening in human Huh7.5 cells validated 23 diverse antiviral drugs. Extending our studies to lung epithelial cells, we found that there are major differences in drug sensitivity and entry pathways used by SARS-CoV-2 in these cells. Entry in lung epithelial Calu-3 cells is pH-independent and requires TMPRSS2, while entry in Vero and Huh7.5 cells requires low pH and triggering by acid-dependent endosomal proteases. Moreover, we found 9 drugs are antiviral in lung cells, 7 of which have been tested in humans, and 3 are FDA approved including Cyclosporine which we found is targeting Cyclophilin rather than Calcineurin for its antiviral activity. These antivirals reveal essential host targets and have the potential for rapid clinical implementation.

## Introduction

Coronaviruses represent a large group of medically relevant viruses were historically associated with the common cold. However, in recent years, members of the coronavirus family have emerged from animal reservoirs into humans and have caused novel diseases (1). First, Severe Acute Respiratory Syndrome (SARS-CoV) emerged in China in 2003, followed by Middle East Respiratory Syndrome (MERS-CoV) in 2012 (2, 3). While SARS was, in the end eradicated, MERS continues to cause infections in the Middle East. Beginning in December 2019, into continuing into January 2020, it became clear that a new respiratory virus was spreading in Wuhan, China. Rapid sequencing efforts revealed a coronavirus closely related to SARS, and was named SARS-CoV-2 (4). Unfortunately, this virus is highly infectious and has spread rapidly around the world.

Identification of broadly acting SARS-CoV-2 antivirals is essential to clinically address SARS-CoV-2 infections. A potential route to candidate antivirals is through the deployment of drugs that show activity against related viruses. Previous studies found that the antiviral drug remdesivir, which was developed against the RNA-dependent RNA polymerase of Ebola virus, was also active against SARS-CoV-2 in vitro, with promising results in clinical trials (5–7). Chloroquine, and its derivatives including hydroxychloroquine are approved for use in malaria, and many in vitro studies have found that these drugs are also active against coronaviruses, including SARS-CoV-2 (8, 9). This led to early adoption of these agents to treat COVID-19 (the disease caused by SARS-CoV-2 infection); however, little efficacy of these agents has been demonstrated in subsequent clinical trials (10). It remains unclear why these agents have not been more active in humans.

There are currently more than 3000 FDA approved drugs, and many others that have been tested in humans. We created an in-house library of 3059 drugs including ~1000 FDA approved drugs and ~2100 drug-like molecules against defined molecular targets with validated pharmacological activity. In addition, we purchased drugs with reported anti-SARS-CoV-2 activity (e.g. remdesivir). Viruses encode unique proteins essential for infection, and most approved antivirals target these virally encoded essential targets. This class of antivirals has been termed direct-acting antivirals. Viruses are also dependent on host cellular machineries for successful infection, and drugs that block these activities are host-targeted antivirals. Given our dearth of effective treatments, we developed a screening platform that would allow us to identify both direct-acting and host-targeted antivirals that can be potentially repurposed for use against SARS-CoV-2 (11).

We developed a specific and sensitive assay to quantify viral infection using a cell-based high-content approach. We began our studies in African green monkey (*Cercopithecus aethiops)* kidney epithelial cells (Vero) because they are routinely used to propagate SARS-CoV-2. They are robustly infected, and thus Vero cells are widely used as a model system to screen for antivirals (8, 12–14). We screened our in-house repurposing library, identifying only six drugs that were antiviral with low toxicity in the primary screen. Given how few candidates emerged, we reasoned that human cells might be a better model of infection and thus tested a panel of human cell lines to identify cells that are easy to grow, and permissive to infection. We found that the human hepatocyte cell line Huh7.5 was readily infected with SARS-CoV-2. Screening in this human cell line we validated 23 drugs that were active in dose-response experiments and showed a favorable selective index versus toxicity (15). These candidates targeted a wide variety of cellular activities, but few were active in Vero cells. However, one class, the chloroquines and their derivatives were active in both cell types.

The entry pathway of SARS-CoV-2 has only begun to be elucidated with much of what we know being inferred from studies of the related SARS-CoV-1(16, 17). The coronavirus glycoprotein, or Spike, requires proteolytic processing for entry (16, 18, 19). This processing can occur outside the cell, or within the endolysosomal compartment (16, 19). Both SARS-CoV-1 and −2 engage angiotensin-converting enzyme 2 (ACE2) as their plasma membrane receptor (20–22). Upon binding, the viruses along with the receptor are endocytosed into the cell into a low pH endosomal compartment where there are proteases, including cathepsins, that can cleave Spike and allow for entry into the cytosol (23–25). Since cathepsins require a low pH for activity, chloroquine and its derivatives that neutralize this low pH can effectively block viral entry (23–25). Recent studies have also identified a plasma membrane-associated serine protease, TMPRSS2, is active against Spike, cleaving the protein extracellularly thereby bypassing the requirement for endosomal proteases (26–28). Whether SARS-CoV-2 enters through different routes in different cell types remains unclear.

Lung epithelial cells are the major cellular target for SARS-CoV-2 in vivo and have been used to explore the role of TMPRSS2 in infection. Perhaps surprisingly, while we found remdesivir was antiviral in Calu-3, hydroxychloroquine was not. Since a panel of quinolines had no activity in Calu-3 cells, these data suggest that entry in lung epithelial cells is independent of low-pH processing in the endosomal compartment. In contrast, the TMPRSS2 inhibitor camostat was highly active in Calu-3 cells but inactive in Vero and Huh7.5 cells. These data demonstrate distinct modes of entry in lung cells (21). Further, these data suggest that there may be other fundamentally different cellular requirements in different cell types. We screened our 23 validated candidates in Calu-3 cells and found only 9 drugs showed activity, including 3 FDA approved drugs: Cyclosporine, Dacomitinib and Salinomycin. In additional studies, we found that cyclosporine analogs that target Cyclophilin A were active against SARS-CoV-2, but not compounds that target Calcineurin. Identifying broadly acting antivirals is essential to move forward with clinical treatments for SARS-CoV-2.

## Results

### Vero cells are permissive to infection and can be used for antiviral screening for direct acting antivirals

SARS-CoV-2 is routinely propagated in Vero E6 cells (12, 14, 26). When growing the virus in either Vero E6 or Vero CCL81 cells, two different strains of Vero cells from ATCC, we observed that SARS-CoV-2 is cytopathic in Vero E6, but not in Vero CCL81 (data not shown) (12). Moreover, viral stocks propagated from either of these cells produced similar titers of virus (1×10^7^ PFU/mL) suggesting that viral replication and cytotoxicity are separable. Therefore, we set out to develop a quantitative microscopy-based assay to measure the level of replication of SARS-CoV-2 more directly in infected cells, and we chose Vero CCL81 to uncouple toxicity from infection. We first validated that our antibodies could detect infection of SARS-CoV-2. We used an antibody to dsRNA, and to SARS-CoV-2 Spike (Figure 1a) (29) (30).

**Figure 1:**
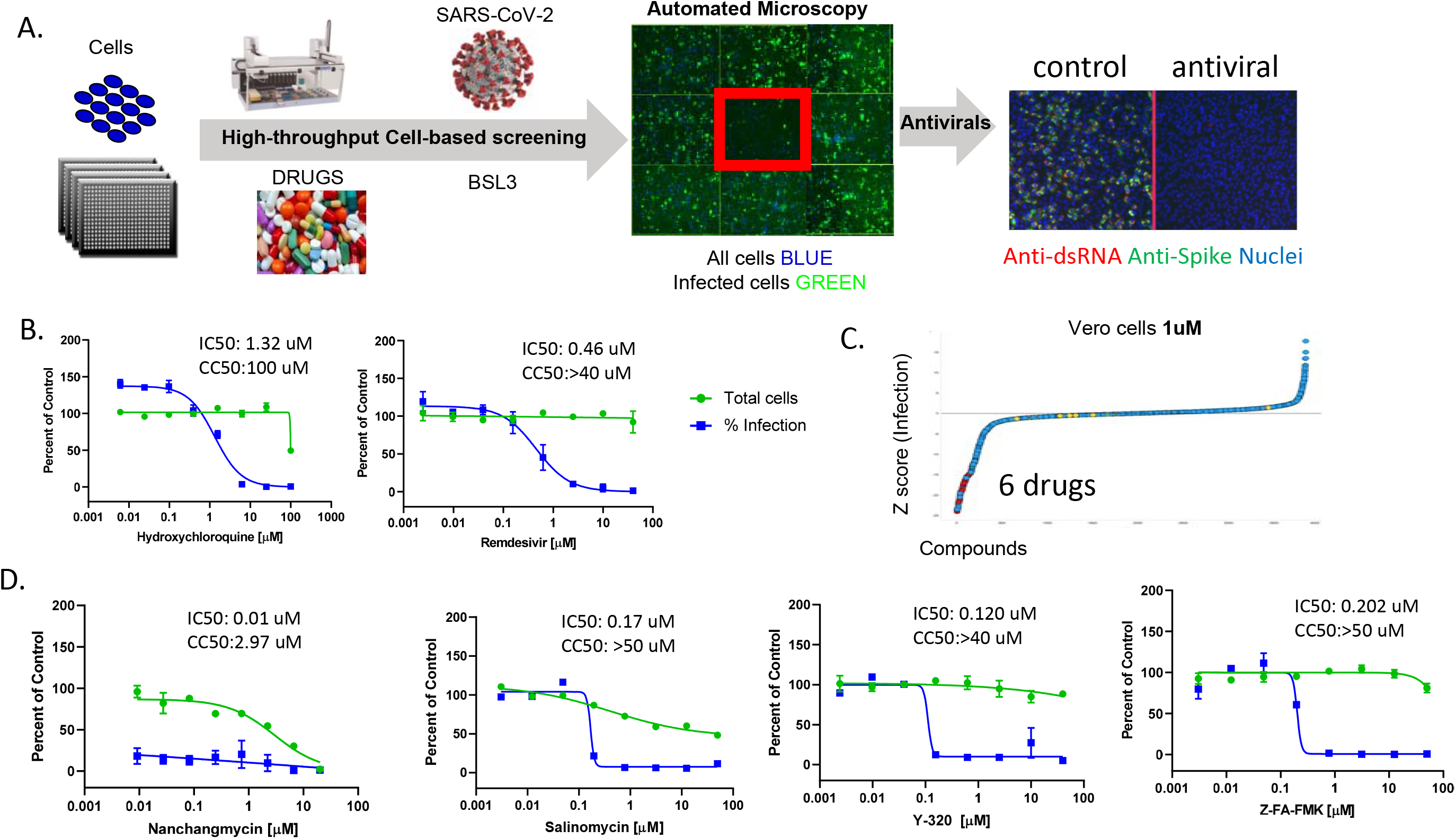
High-throughput screening in Vero cells to identify antivirals against SARS-CoV-2. A. Schematic of the screening strategy. Vero cells were plated in 384 well plates, drugs were added and the cells were infected with SARS-CoV-2 (MOI=1). 30 hpi cells were stained for viral infection (dsRNA, Spike) and imaged using automated microscopy to define cell number and percent infection. Antivirals show little impact on cell number and block viral infection. B. Dose-response analysis of Vero cells treated with hydroxychloroquine or remdesvir. C. Z-scores of the Vero drug screen performed at 1μM. 6 drugs had >60% reduction in infection with >80% cell viability. D. Dose-response analysis of three candidates identified in the screen.

We created an in-house library of 3059 drugs purchased from Selleckchem. This library contains ~1000 FDA approved drugs and ~2100 drug-like molecules against defined molecular targets with validated pharmacological activity. The library contains 678 known kinase inhibitors, 435 annotated cancer therapeutics, 190 epigenetic regulators, 411 anti-viral/infectives, 596 GPCR and ion channel regulators. The remaining compounds falling into diverse target classes. We next optimized the dose and timing of infection by performing dose-response studies with known antivirals. Indeed, we found that hydroxychloroquine and remdesivir were active in Vero cells and presented with little cytotoxicity at the active doses (Figure 1b) (8). Next, we validated the assay metrics, and observed a Z’=0.7 (Figure S1) (31).

We used this assay pipeline to screen our in-house repurposing library in 384 well plates at a final concentration of 1 μM (Figure 1c) (32). We quantified the percentage of infected cells as well as the total cell number per well, to allow for exclusion of toxic compounds. We robustly identified the positive control remdesivir as antiviral (Figure S1) (8). Using a threshold of <40% infection and >80% viability, as compared to the vehicle control, we identified only six drugs that were antiviral in our primary screen (Table S1). This included the natural product nanchangmycin, which we previously found in a drug repurposing screen against Zika virus (32). Nanchangmycin was broadly antiviral against viruses that enter cells through endocytosis, consistent with the role of endosomal acidification for SARS-CoV-2 entry in these cells (32). We then repurchased powders and validated four of these candidates in a dose-response assays where we observed antiviral activity in the absence of toxicity (Figure 1d).

### Human hepatocyte Huh7.5 cells are permissive to infection and can be used to identify antivirals

Since Vero cells are derived from African green monkeys, we set out to identify a human cell line permissive to infection. To this end, we infected a panel of human cell lines with SARS-CoV-2 and monitored infection by microscopy. We initially tested A549, Calu-1, Huh7, Huh7.5, HepG2, HaCaT, IMR90, NCI-H292, CFBE41o, and U2OS cells. We detected less than 1% infection of A549, Calu-1, Huh7, HepG2, HaCaT, IMR90, NCI-H292, CFBE41o, and U2OS cells (data not shown). Interestingly, while Huh7 were largely non-permissive, the derivative cell line Huh7.5 cells was permissive to SARS-CoV-2 (Fig 2a). Huh7.5 cells are defective in innate immune signaling (RIG-I) and are known to be more permissive to many viruses, including Hepatitis C virus(15). Remdesivir and hydroxylchloroquine were antiviral against SARS-CoV-2 in Huh7.5 cells with IC50s that were greater than 10-fold lower than those observed in Vero cells (Fig 2b). We also found that nanchangmycin was antiviral against SARS-CoV-2 in Huh7.5 cells (Fig S2).These observations suggest that Huh7.5 cells may be more sensitive to some classes of inhibitors, and may reveal antivirals that are selectively active against human targets.

**Figure 2.**
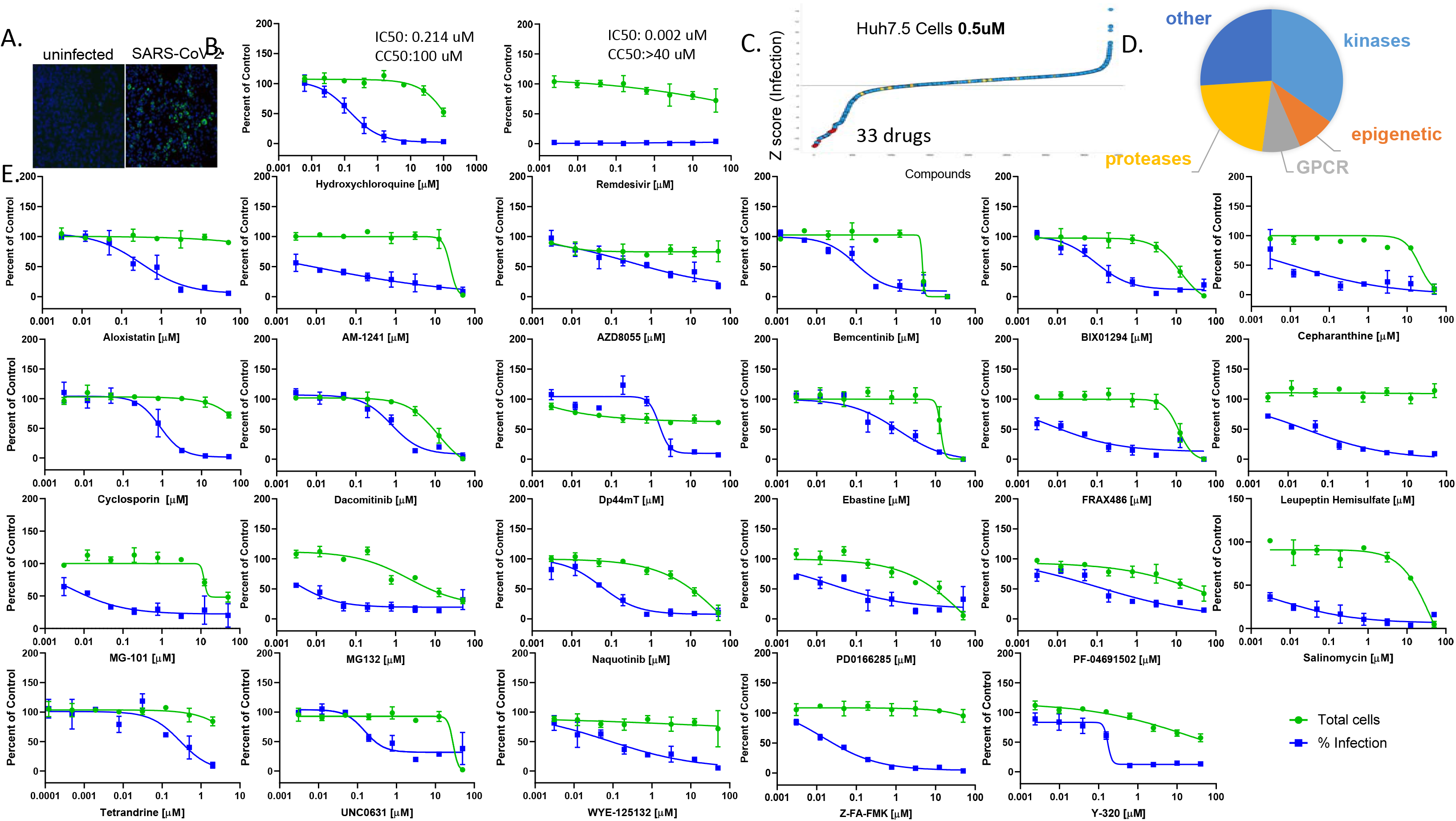
High-throughput screening in human Huh7.5 cells to identify antivirals against SARS-CoV-2. A. Huh7.5 cells were infected with SARS-CoV-2 (MOI=1) and 30hpi processed for microscopy. B. Dose-response analysis of Huh7.5 cells treated with hydroxychloroquine or remdesvir. C. Z-scores of the Huh7.5 drug screen performed at 0.5μM. 33 drugs had >60% reduction in infection with >80% cell viability. D. Dose-response analysis of the candidates with a selective index >2 identified in the screen.

We optimized our image-based assay in Huh7.5 cells using remdesivir and observed a Z’=0.61 (Fig S3) (31). We screened our repurposing library at 500nM quantifying both the percentage of infected cells as well as cell number to exclude toxic compounds (Fig 2c). We found 33 drugs had antiviral activity in the absence of cytotoxicity (<40% infection, >80% viability, as compared to vehicle control) (Table S2). This included three of the six drugs identified in Vero cells: Z-FA-FMK, Y-320 and Salinomycin.

We repurchased powders for 32 drugs and tested their activity in dose-response assays in Huh7.5 cells against SARS-CoV-2. Cell number and the percent of infected cells were quantified. Remdesivir and hydroxychloroquine were used as positive controls and vehicle controls (DMSO) was included as a negative control (8). Of those tested, 23 drugs showed activity and fell into diverse classes (Fig 2d). Dose-response curves are shown for 23 candidates and the IC50s and CC50s were calculated (Fig 2e). The Selectivity Index (SI, ratio between antiviral and cytotoxicity potencies) was calculated and the 23 candidates were antiviral with SI>3 (Figure 2e, Table S3). Dose-responses curves for the other candidates that did not validate in Huh7.5 cells are shown in Fig S4.

Direct-acting antivirals are likely to be active against the virus in multiple cell types, as was observed for remdesivir. In addition, host-directed antivirals that target key steps in the viral lifecycle and are highly conserved and broadly expressed are also likely to emerge across cell types. One example is the endosomal acidification blocker hydroxychloroquine which indeed scored as antiviral in both cell types (9, 23, 24). In total, we identified three drugs as antiviral in both screens. We performed dose-responses in Vero cells against the candidates from the Huh7.5 screen. We found that 5 additional compounds were antiviral in Vero cells with a SI>3, AZD8055, BIX01294, Ebastine, MG-132, and WYE-125132, albeit at higher concentrations (Figure S5). However, the majority of the antivirals that were validated in Huh7.5 cells were not active in Vero cells.

### Lung epithelial cells show differences in drug sensitivities

We next focused on lung epithelial models as these are the most relevant to human infections. We found that a number of lung-derived epithelial cell lines were refractory to infection (eg A549, Calu-1, NCI-H292, CFBE41o). However, we found that Calu-3 cells, that have been shown to be permissive for many coronaviruses including SARS-CoV-2, were highly readily infected (Fig 3a) (14, 26, 33). We optimized assays using Calu-3 cells and tested their sensitivity to remdesivir and hydroxycholorquine. As expected, we found that while the direct acting antiviral remdesivir was antiviral; however, hydroxychloroquine had little or no activity in Calu-3 cells (Fig 3b). This led us to test the antiviral activity of a panel of chloroquine derivatives and we found that none of these had activity against SARS-CoV-2 (Fig 3c), while these compounds are antiviral in both Vero cells and Huh7.5 cells (Fig 3d). This suggests that there are major differences in the requirement for endosomal acidification during infection of SARS-CoV-2 in lung epithelial cells.

**Figure 3.**
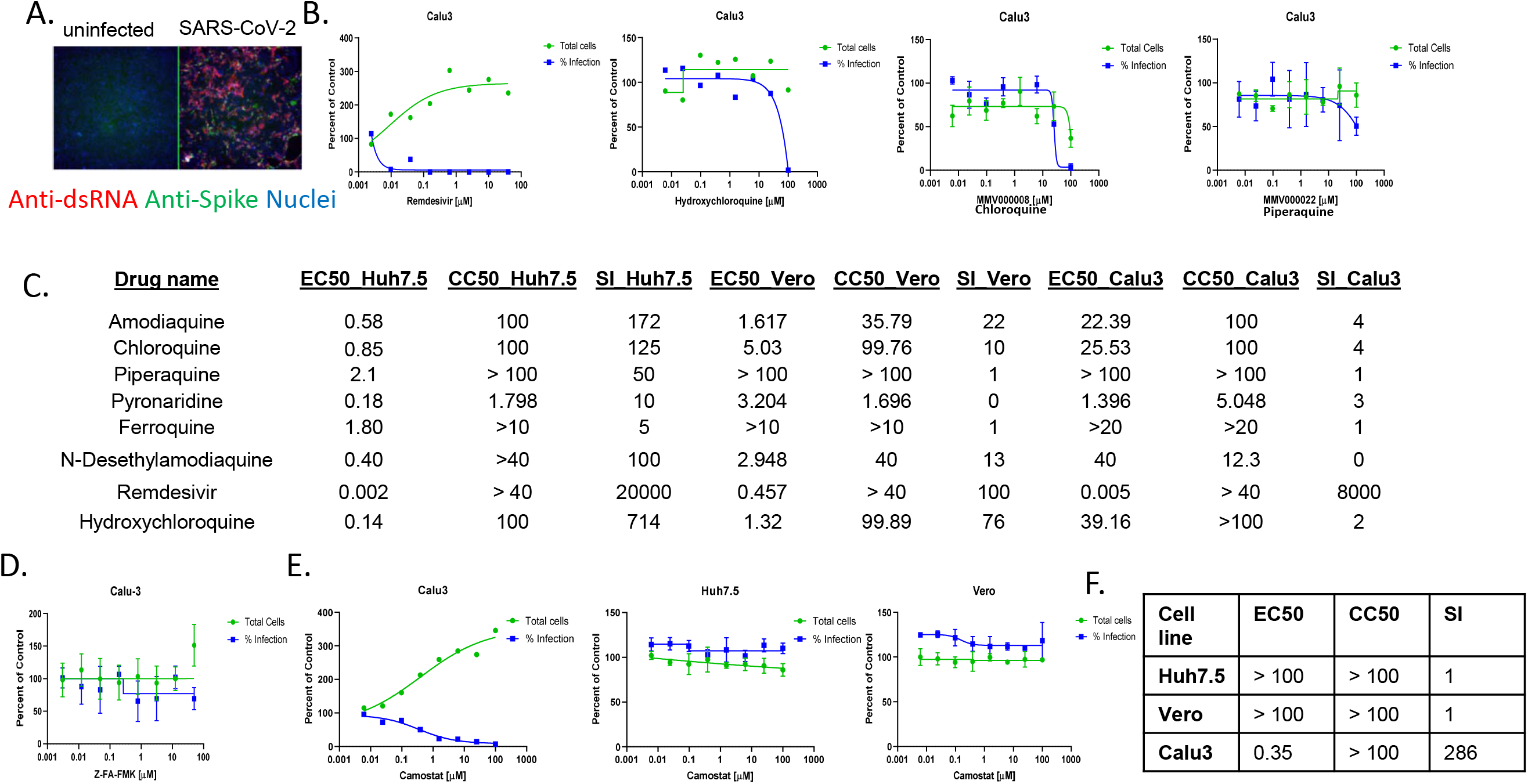
Cell type specific dependencies of entry inhibitors. A. The human lung epithelial Calu-3 cells were infected with SARS-CoV-2 (MOI=0.5) and 48 hpi processed for microscopy. B. Dose response analysis of Calu-3 cells treated with quinolines or remdesvir. C. IC50, CC50 and SI for Vero, Huh7.5 and Calu-3 cells treated with a panel of quinolines or remdesivir. D. Dose response analysis of Calu-3 cells treated with cathepsin inhibitor Z-FA-FMK. E. Dose response analysis of Calu-3, Vero and Huh7.5 cells treated with camostat. F. IC50, CC50 and SI for camostat across cell types.

Endosomal acidification is thought to be required for SARS-CoV-2 entry to maintain the low pH necessary for endosomal cysteine protease activity required for priming Spike for membrane fusion (26). Consistent with the requirement for acidification in Vero and Huh7.5, the cathepsin inhibitor Z-FA-FMK emerged as antiviral in both cell types (Fig 1d, Fig 2e). We tested Z-FA-FMK in Calu-3 cells and found that it had no antiviral activity (Fig 3d), consistent with a lack of a requirement for endosomal acidification. Recent studies found the plasma membrane-associated serine protease, TMPRSS2, can prime the viral glycoprotein for entry in lung epithelial cells (26). Therefore, we tested the role of TMPRSS2 by treating cells with the inhibitor camostat. We found that camostat was antiviral in Calu-3 cells but had no activity in either Vero or Huh7.5 cells (Figure 3e-f) (26). Moreover, the main endosomal kinase Phosphatidylinositiol-3-Phosphate/Phosphatidylinositol 5- kinase, PIKfyve, promotes internalization of diverse viruses and was recently shown to impact entry of coronaviruses including SARS-CoV-2 in HeLa cells (34). Using the PIKfyve inhibitor apilimod, we found that PIKfyve promotes infection of SARS-CoV-2 in Huh7.5 and Vero cells, with little importance Calu-3 cells (Fig S6). These data suggest that the entry pathway used by SARS-CoV-2 is cell-type specific.

### Nine candidates are antiviral against SARS-CoV-2 in lung epithelial cells

To determine which of the 23 antiviral candidates validated in Huh7.5 cells also had antiviral activity in Calu-3 cells we performed dose-response studies. We found that 9 drugs were antiviral against SARS-CoV-2 in Calu-3 cells with a selectivity index greater than 2 (Fig 4). These include: two drugs with unclear targets (Salinomycin, Y-320), kinase inhibitors (AZD8055, bemcentinib, dacomitinib, WYE-125132), histamine receptor inhibitor (ebastine), iron chelator Dp44mT, and the cyclophilin inhibitor cyclosporine. Many kinase inhibitors were quite potent, suggesting an important role in intracellular signaling for infection. The other drugs tested in Calu-3 with a SI<3 are shown in Fig S7. The full table of candidates from the Huh7.5 screen with IC50, CC50 and SI are shown in Figure S8.

**Figure 4.**
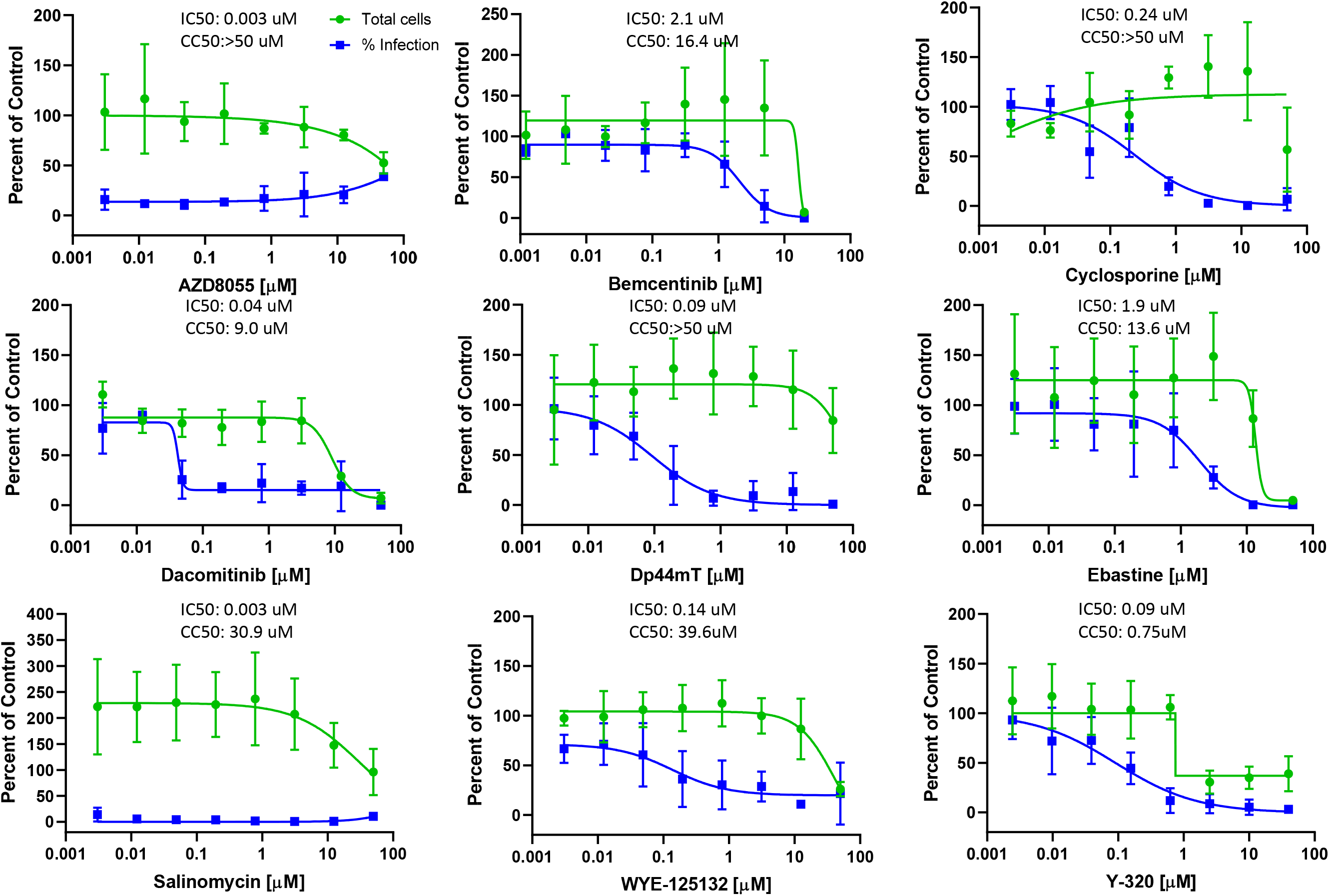
Validation of antiviral activity of 9 candidates in Calu-3 cells. Dose-response analysis of the Huh7.5 candidates in Calu-3 cells with a selective index >2.

### Cyclosporine is antiviral likely through interactions with cyclophilins

Cyclosporine is an FDA approved generic drug that is readily available and showed a sub-micromolar IC50 with high selectivity in both Huh7.5 and Calu-3 cells (Fig 3, Fig 4, Fig S8). Cyclosporine binds Cyclophilin A and prevents activation of the phosphatase Calcineurin which is required for the nuclear translocation of the nuclear factor of activated T cells (NFAT) (35–37). Inhibition of this pathway in T cells is used as an immunosuppressant (38). Cyclosporins have been shown to have antiviral activity against a wide variety of viruses, including other coronaviruses (39–52). The activity of Cyclosporine against previously studied coronaviruses is Cyclophilin-dependent and independent of Calcineurin (50, 51). We set out to perform initial structure-activity relationships (SAR) and to determine if this activity was through its inhibition of Cyclophilin or inhibition of Calcineurin. For these studies, we obtained a panel of Cyclosporine analogs including Cyclosporin A, Cyclosporin B, Cyclosporin C, Cyclosporin H and Isocyclosoporin A (53). We found that Isocyclosporin A, Cyclosporin A, Cyclosporin B, and Cyclosporin C were active with increasing IC50s (Fig 5a-c). Cyclosporine H, shows only weak binding to Cyclophilin A, and has no immunosuppressant activity, as it does not inhibit the phosphatase activity of Calcineurin (53, 54). Cyclosporin H has a log reduction in activity compared to Cyclosporine, but retains some activity. PSC 833 is a non-immunosuppressant derivative of cyclosporine that does not inhibit Calcineurin, has a similar activity to Cyclosporin C(55). TMN355 is a Cyclophilin A inhibitor that is 27 times more potent than Cyclosporine A in inhibiting the prolyl isomerase activity of cyclophilin A (56). We found the latter to lack antiviral activity, suggesting that the enzymatic activity of Cyclophilin A is dispensable.

**Figure 5.**
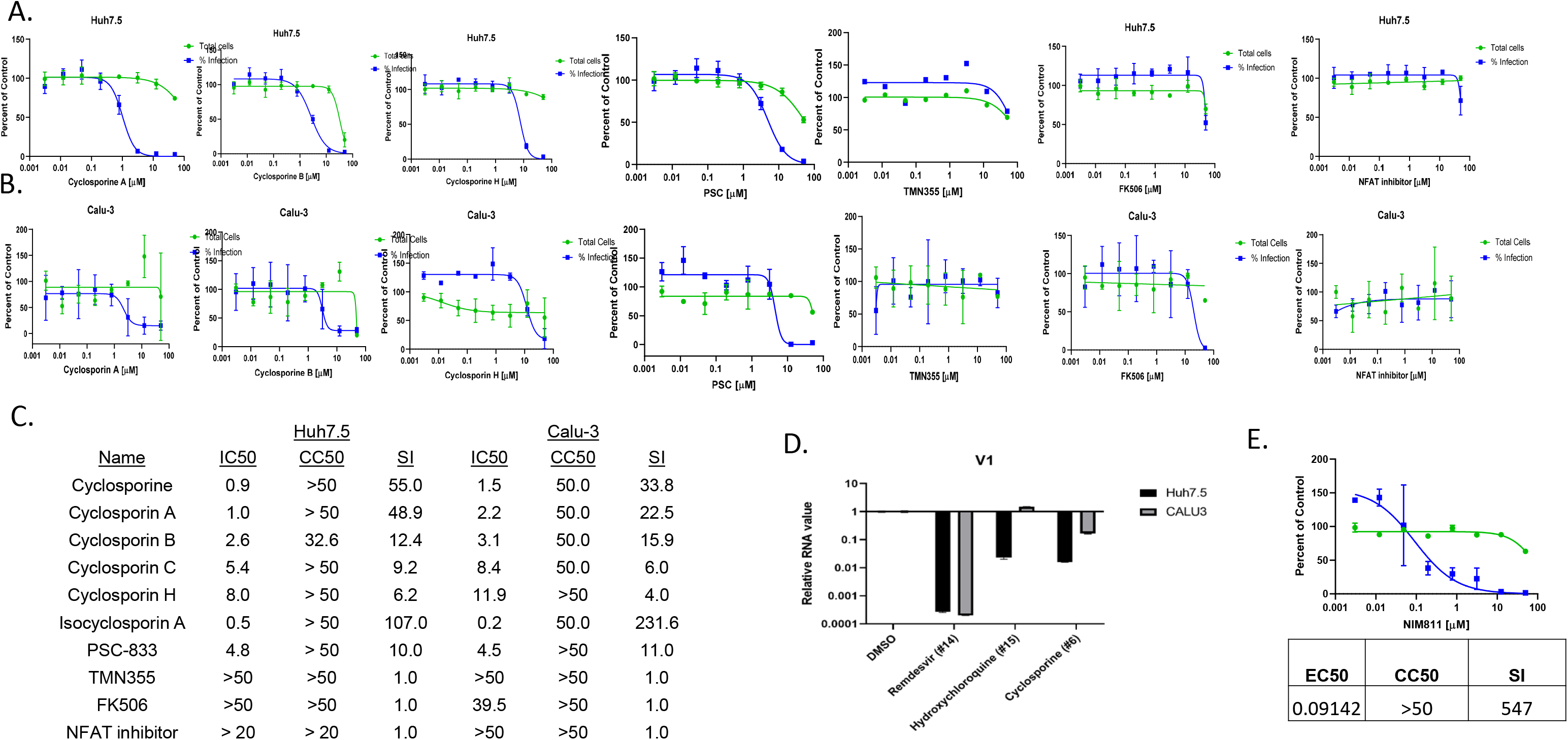
Cyclosporine is antiviral against SARS-CoV-2 independent of calcineurin. A. Dose-response analysis of Huh7.5 cells treated with a panel of cyclosporins and related drugs. B. Dose-response analysis of Calu-3 cells treated with a panel of cyclosporins and related drugs. C. Table of IC50s, CC50s and SI of Huh7.5 cells and Calu-3 cells treated with the indicated drugs. D. RT-qPCR analysis of viral replication in Huh7.5 and Calu-3 cells treated with the indicated drugs. Mean±SE shown. E. Dose-response analysis of Huh7.5 cells treated with NIM811. Table of IC50, CC50 and SI shown.

We further validated that cyclosporine is antiviral in both cell types by performing RT-qPCR (Fig 5d). In addition, NIM811 is another non-immunosuppressive cyclosporine derivative of cyclosporine, and we found that it is potently antiviral (Fig 5e), further suggesting that the antiviral activity is Cyclophilin-dependent and separable from Calcineurin. Strikingly, the activity of this panel of drugs are similar in the two cell lines. These data suggest that Cyclosporine has the same target and mechanism-of-action. We also found that none of these drugs are antiviral in Vero cells (Figure S9).

To further assess the mechanism by which Cyclosporine is antiviral, we tested FK506, an inhibitor of Calcineurin. FK506 binds the related immunophilin FKBP, rather than Cyclophilin A, to block the phosphatase activity of Calcineurin, and thus is also a potent immunosuppressant (57). We found that FK506 has no activity against SARS-CoV-2 (Fig 5a-c). Moreover, since one of the major targets of Calcineurin is the activation of NFAT, we also tested whether an NFAT inhibitor impacted viral infection (58). We found that the NFAT inhibitor had no effect on infection (Fig 5a-c). Altogether, we found that cyclosporins are potent antivirals against SARS-CoV-2 in lung epithelial cells, and that this activity is independent of Calcineurin and NFAT.

## Discussion

The emergence of SARS-CoV-2 has led to devastating global morbidity and mortality, creating an immediate need for new therapeutics and vaccines. Repurposing existing drugs can allow for rapid deployment of therapeutics that have already been tested in humans (59). Remdesivir was developed against the Ebola virus RNA-dependent RNA polymerase, and was also found to have robust activity against SARS-CoV-2 (8). Importantly, we found that remdesivir is active against SARS-CoV-2 across cell types. Chloroquine and hydroxychlorquine have been used for decades to treat malaria and have been shown to have in vitro antiviral activity against SARS-CoV-2 (8, 9). However, we find that this antiviral activity is cell-type specific. Lung epithelial cells are resistant to these drugs, and this may explain the lack of efficacy seen in many trials (10).

To determine if there are additional drugs that are active against SARS-CoV-2 in vitro, we screened a repurposing library that includes ~1000 FDA approved drugs and ~1000 additional drugs that have been tested in humans. Repurposing can be used to reveal new and similar pathways and targets, but also the time and monetary investment associated with repurposing is potentially less since these drugs often bypass Phase-1 trials (60, 61). Initial screens in Vero cells yielded few active drugs, leading us to pursue a screen in human Huh7.5 cells, a transformed hepatocyte line deficient in innate immune signaling. Using this model system, we identified 33 drugs, and validated 23 with dose-response assays. This includes many drugs that were previously shown to have activity against other coronaviruses (tetrandrine, cepharanthine, cyclosporine, alixostatin, MG132, salinomycin), and SARS-CoV-2 (salinomycin, tetrandrine, cepharanthine, cyclosporine, ebastine) (13, 40, 50, 54, 62–68).

The 23 drugs fall into distinct classes and most have known targets. However, two drugs that were active across cell types, Salinomycin and Y-320, do not have clear targets. Salinomycin, is a polyether antibiotic and chemotherapy drug, that has been shown to be antiviral against many viruses, including coronaviruses (13, 68–70). Salinomycin was also identified in a Vero cell screen (13). Mechanistically, some studies have suggested that Salinomycin is an ionophore that can attenuate viral entry by disrupting the acidification of the endosome (71). Other studies have implicated Salinomycin in ER stress (72). Studies in mice have shown antiviral activity against influenza (71). Salinomycin has also been characterized as an activator of autophagy, which may influence SARS-CoV-2 infection (73, 74) (75). Y-320 is a phenylpyrazoleanilide immunomodulatory agent that has been shown to inhibit IL-17 production by T cells and has activity in monkeys (76). Interestingly, treatment with Y-320 is associated with decreased IL-6 production, a cytokine that is thought to be highly expressed in SARS-CoV-2 infection (76–78). However, it is unclear how Y-320 could attenuate SARS-CoV-2 in non-immune cells.

Ebastine is a potent H1-histamine receptor antagonist, used for allergic disorders outside of the US, particularly in Asia (79). We found that ebastine is antiviral in all three cell types, although 10-fold less active in Vero cells (13). Ebastine is orally available with few side effects and there are there clinical trials underway in China testing whether ebastine can impact COVID-19 outcomes (80). Since other H1-histamine receptor antagonists were not active, it is unclear why this particular agent is more effective at inhibiting SARS-CoV-2 infection. Interestingly, ebastine and its active metabolite, carebastine, are reported to inhibit expression of IL-6, while many other H1-histamine receptor antagonists do not (81, 82).

We also identified 6 protease inhibitors as antiviral in Huh7.5 cells. Two cysteine protease inhibitors, Z-FA-FMK and MG-132, had activity in both Vero and Huh7.5 cells. None of the protease inhibitors were active in Calu-3 cells. This observation suggests that they are not targeting the viral proteases. Consistent with this, Z-FA-FMK is an inhibitor of cathepsins which are required for SARS-CoV-2 entry in cells where endosomal proteases are required for Spike cleavage, and thus we observe no requirement in Calu-3 cells where TMPRSS2 is required for infection (16, 19, 26–28).This has important implications in diverse SARS-CoV-2 studies where there may be cell-type specific requirements for different steps in the replication cycle.

We also identified two inhibitors against the cellular histone methyltransferase G9a as antiviral in Huh7.5 cells. However, these drugs were not active in Calu-3 cells, suggesting that there are cell type specific requirements. AM1241 is a selective cannabinoid CB2 receptor agonist that we found was antiviral in Huh7.5 cells. GW842166X, another CB2 agonist that has a 10-fold higher EC50, was not active. Moreover, dose-response studied found that AM1241 is not active in either Vero or Calu-3 cells.

Cepharanthine and tetrandrine are both bis-benzylisoquinoline alkaloids produced as natural products from herbal plants (83). Tetrandrine, a traditional Chinese medicine and calcium channel blocker, has been shown to antagonize calmodulin. It has anti-tumor and anti-inflammatory effects, and can effectively inhibit fibroblasts, thereby inhibiting pulmonary fibrosis (84, 85). Multiple studies have suggested that tetrandrine has antiviral activity, including against dengue virus and herpes simplex 1 virus (86, 87). Tetrandrine has also been shown to inhibit entry of Ebola virus into host cells *in vitro* and showed therapeutic efficacy against Ebola in preliminary studies on mice (88). Currently, there is an ongoing clinical trial using tetrandrine in COVID-19 patients to improve pulmonary function (80). Cepharanthine is reported to have anti-inflammatory and immunoregulatory properties and is used to treat a variety of acute and chronic conditions outside of the US (89). Both cepharanthine and tetrandrine were previously shown to have antiviral activity against the human coronavirus OC43 and in recent studies on SARS-CoV-2 in Vero cell screens (13, 62, 63). While both of these molecules were antiviral in our Huh7.5 screen, neither were active in Calu-3 cells. This may suggest that they are modulating endosomal entry pathways.

We identified few metabolic regulators. Dp44mT is a potent iron chelator that we found to be antiviral against SARS-CoV-2 in Huh 7.5 and Calu-3 cells (56). A clinical trial with the iron chelator deferoxamine is underway (NCT04333550). However, other iron chelators in our library, deferasirox and deferiprone, were not identified as antiviral making the mechanism of action unclear.

We identified several kinase inhibitors as antivirals against SARS-CoV-2. FRAX486 is a p21-activated kinase (PAK) inhibitor that is antiviral in Huh7.5 cells, but only modestly impacted infection of Calu-3 cells (90). Other PAK inhibitors were not identified in our screens. PAK is required for entry by many viruses(91). PD0166285 is a potent Wee1 and Chk1 inhibitor that is antiviral in Huh7.5 cells, but shows strong toxicity in Calu-3 cells (92).

We also found three mTOR inhibitors, AZD8055, PF-04691502, and WYE-125132 are antiviral against SARS-CoV-2 in Huh-7 and Calu-3 cells. These are highly potent ATP competitive mTOR inhibitors that target both TORC1 and TORC2. In our library, none of the rapamycin analogs that selectively inhibit mTORC1 were active. We also identified two potent selective and irreversible inhibitors of EGFR, dacomitinib and naquotinib. The other 44 EGFR inhibitors showed no activity in Huh7.5 cells. Importantly, dacomitinib is a potent antiviral in Calu-3 cells. It is unclear if the target is indeed EGFR, but for many viruses EGFR activation promotes viral entry which may also be the case for SARS-CoV-2 (93–97).

Bemcentinib is a first-in-class Axl inhibitor that we found inhibits SARS-CoV-2 infection of Huh7.5 cells and Calu-3 cells (98). Axl can be used as an attachment factor for the entry for many viruses including Ebola and Zika virus (99, 100). While not published, news releases suggest that Bemcentinib has demonstrated promise in preclinical data against early infection with the SARS-CoV-2. A fast-tracked clinical trial is underway in the UK (101).

Cyclosporine is a commonly used immunosuppressant that binds Cyclophilin A and inhibits the calcium-dependent phosphatase Calcineurin which is required for the nuclear translocation of the nuclear factor of activated T cells (NFAT) (35–38, 41). Inhibition of this pathway in T cells is used as an immunosuppressant. We found that Cyclosporine is active in both Huh7.5 and Calu-3 cells, but has no activity in Vero cells nor did the cyclosporine analogs. A recent screen in Vero cells did find activity with Cyclosporine against SARS-CoV-2 (13). Cyclophilin A is a ubiquitously expressed peptidyl-prolyl *cis-trans* isomerase (102). Cyclophilin A and other Cyclophilins have chaperone-like activity and take part in protein-folding processes (103). Cyclophilin A has been shown to be an important cellular factor that facilitated many diverse viral infections. This includes human immunodeficiency virus type 1 (HIV-1), influenza virus, hepatitis C virus (HCV), hepatitis B virus (HBV), vesicular stomatitis virus (VSV), vaccinia virus (VV), severe acute respiratory syndrome coronavirus (SARS-CoV) and Rotavirus (RV) (41–49, 104, 105). The coronaviruses HCoV-229E, HCoV-NL63, FPIV, mouse hepatitis virus (MHV), avian infectious bronchitis virus, and SARS-CoV have been found to be attenuated by Cyclosporin A (40, 64, 106). Cyclosporine and its non-immunosuppressive derivatives can inhibit replication of a number of viruses including some coronaviruses. In most cases the responsible cyclophilin is CypA (54, 105), but CypA and CypB were found to be required for FCoV replication (106). For HCoV-NL63, and HCoV-229E, cyclophilin A is required for infection in CaCo-2 cells (50) and Huh-7.5 cells respectively (51, 52). It is generally thought that the activity of Cyclosporine against coronaviruses is Cyclophilin-dependent and independent of Calcineurin.

We found that a number of Cyclosporins were antiviral with similar potencies including Cyclosporine, Cyclosporin A, Cyclosporin B and the metabolic breakdown product of Cyclosporin A, Isoscyclosporin A. We also found that Cyclophilin A is likely required, as Cyclosporin H, which is a weak binder had reduced activity. However, the enzymatic activity of Cyclophilin A is likely dispensable as TMN355 was inactive. To further address the role of Calcineurin, we tested a non-immunosuppressant derivative of cyclosporine that does not inhibit calcineurin, has a similar activity to Cyclosporin C. We also found that FK506, a Calcineurin inhibitor independent of Cyclophilin A, and NFAT inhibitors also have no antiviral activity. Altogether, we found that cyclosporins are potent antivirals against SARS-CoV-2 in lung epithelial cells, and that this activity is independent of calcineurins. NIM811 is a Cyclophilin inhibitor independent of Calcineurin, and we found that this is highly active in Huh7.5 cells, further suggesting that Cyclophilin is required for SARS-CoVo-2 infection. Strikingly, the activities of all of these drugs is similar in the two cell lines suggesting the same target and mechanism-of-action and that Cyclosporine would block SARS-CoV-2 in diverse infected tissues in vivo.

One approach would be to use Cyclophilin inhibitors that do not have immunosuppressive activity such as NIM811 or others that have been tested for HCV infection (Alisporivir (Debio-025) and SCY-635) (107) or for HIV infection (NIM818) (108) (54). Another possibility is to use Cyclophilin inhibitors that also target Calcineurin (eg. Cyclosporine). One of the major complications of COVID-19 is the hyper-inflammatory response and cytokine storm associated with increased immune activation. To prevent hyper-activation, there has been interest in treating COVID-19 patients with immunosuppressants (109). There are ongoing trials for a variety of agents including anti-IL6 and JAK inhibitors, two clinical trials using sirolimus, the FDA approved mTOR inhibitor, which selectively inhibits mTORC1. We find no antiviral activity of sirolimus or other rapamycin derivatives. In contrast, Cyclosporin A is an approved immunosuppressant that we found is also antiviral at concentrations achieved in vivo (110). Therefore, it may be useful to implement clinical trials using Cyclosporin A as an immunosuppressant as it would potentially ameliorate symptoms by two mechanisms (111).

There have been a large number of screens posted in the literature that suggest antiviral activity of several existing drugs (e.g. azithromycin, Faviprivir, Lopinavir, ribavirin, and ritonavir, tetracycline, etc). These drugs and most screens have been performed in Vero cells, with toxicity as read-outs. Medicines for Malaria Venture (MMV) has compiled a list of drugs that has support for antiviral activity against SARS-CoV-2 (https://www.mmv.org/mmv-open/covid-box). We tested >60 of the 80 compounds and find that in addition to the quinolines and drugs found in our screen there are few additional compounds that show activity at less than 2 uM. While it is possible that some of these drugs are false negatives in our screens, it is likely that many of these candidates do not have antiviral activity when either measuring viral antigen production or when looking in different cell types. It is very important that identified antivirals be tested for their impact on viral replication more directly. Moreover, given the striking differences in sensitivities across cell types it is important to validate the activity of any new antivirals in lung epithelial cells.

Altogether, these studies highlight the roles of cellular genes in viral infection, cell type differences, and our discovery of nine broadly active antivirals suggest new avenues for therapeutic interventions. We found that of the 9 drugs are antiviral in lung epithelial cells, 7 have been used in humans, 3 of these are FDA approved in the US (Cyclosporine, Dacomitinib, and Salinomycin), and Ebastine is approved outside of the US. While clinical trials are underway with some of these candidates, additional trials will be needed to determine the efficacy of these antivirals in COVID-19 patients, to inform future treatment strategies.

## Methods

### Viruses and cells

Vero E6 cells and Vero CCL81 were obtained from ATCC and were cultured in DMEM, supplemented with 10% (v/v) fetal bovine serum, 1% (v/v) penicillin/streptomycin, 1% (v/v) L-Glutamax and were maintained at 37°C and 5% CO2. Huh7.5 cells were obtained from C. Rice (Rockefeller) and cultured in DMEM, supplemented with 10% (v/v) fetal bovine serum, 1% (v/v) penicillin/streptomycin, 1% (v/v) L-glutamine, and were maintained at 37°C and 5% CO2. Calu-3 cells (HTB-55) were obtained from ATCC and cultured in MEM, supplemented with 10% (v/v) fetal bovine serum, 1% (v/v) penicillin/streptomycin, 1% (v/v) L-glutamine, and were maintained at 37°C and 5% CO2. SARS-CoV-2 was obtained from BEI (WA-1 strain). Stocks were prepared by infection of Vero E6 cells in 2% serum plus 10mM HEPES for five days, freeze-thawed, and clarified by centrifugation (PO). Titer of stock was determined by plaque assay using Vero E6 cells and were 1×10^7^ pfu/mL and 1.5×10^6^ TCID50/mL (6). This seed stock was amplified in Vero CCL81 (P1) at 1.5×10^6^ TCID50/mL. All work with infectious virus was performed in a Biosafety Level 3 laboratory and approved by the Institutional Biosafety Committee and Environmental Health and Safety.

### Infections

Cells were plated in 384 well plates (20μL/well) 3,000 cells per well for Vero, 3,000 cells per well Huh7.5, 7,500 cells per well Calu-3. The next day, 50nL of drugs were added. The positive control remdesivir and the negative control DMSO were spotted on each plate. One hour later cells were infected with SARS-CoV-2 (Vero, MOI=1; Huh7.5 MOI=1; Calu-3 MOI=0.5) Cells were fixed (30hpi Vero and Huh7,5, 48hpi Calu-3) in 4% formaldehyde/PBS for 15min at room temperature and then washed three times with PBST. Cells were blocked (2% BSA/PBST) for 60 minutes and incubated in primary antibody (anti-dsRNA J2) overnight at 4C. Cells were washed 3x in PBST and incubated in secondary (anti-mouse alexa 488 and hoescht 33342) for 1h at room temperature. Cells were washed 3x in PBST and imaged using ImagXpress Micro using a 10X objective. Four sites per well were captured. The total number of cells and the number of infected cells were measured using MetaXpress 5.3.3 cell scoring module, and the percentage of infected cells was calculated. The aggregated infection of the DMSO and remdesivir control wells (n=16) on each assay plate were used to calculate z’-factors, as a measure of assay performance and data quality. Sample well infection was normalized to aggregated DMSO plate control wells and expressed as Percentage of Control [POC = (%Infection_sample_/ Average %Infection_DMSO_)*100] and Z-score [Z= (%Infection_sample_ – Average %Infection_DMSO_) / Standard Deviation %Infection_DMSO_]in Spotfire (PerkinElmer). Candidate hits were selected as compounds with POC<40% and viability >80%, compared to vehicle control.

Candidate drugs were repurchased as powders from Selleckchem, MedchemExpress, and MedKoo and suspended in DMSO. Drugs were arrayed in 8-pt dose-response in 384 well plates. Infections were performed using screening conditions. DMSO (n=32) and 10 μM remdesivir (n=16) were included on each validation plate as controls for normalization. Infection at each drug concentration was normalized to aggregated DMSO plate control wells and expressed as percentage-of-control (POC=% Infection _sample_/Avg % Infection _DMSO cont_).). A non-linear regression curve fit analysis (GraphPad Prism 8) was performed on POC Infection and cell viability using log10 transformed concentration values to calculate IC50 values for Infection and CC50 values for cell viability for each drug/cell line combination. Selectivity index (SI) was calculated as a ratio of drug’s CC50 and IC50 values (SI = CC50/IC50).

### RT-qPCR

Huh7.5 (750,000 cells/well) or Calu-3 cells (750,000 cells/well) were plated in 6 well plates. The next day, drugs were added and one hour later infected with SARS-CoV-2 (MOI=0.5). Total RNA was purified using Trizol (Invitrogen) followed by RNA Clean and Concentrate kit (Zymo Researc) 24 hpi for Huh7.5 or 48 hpi for Calu-3. For cDNA synthesis, reverse transcription was performed with random hexamers and Moloney murine leukemia virus (M-MLV) reverse transcriptase (Invitrogen). Synthesized RNA was used as a standard (BEI). Gene specific primers to SARS-CoV-2 (Wuhan v1, NSP14) and SYBR green master mix (Applied Biosystems) were used to amplify viral RNA and 18S rRNA primers were used to amplify cellular RNA using the QuantStudio 6 Flex RT-PCR system (Applied Biosystems). Relative quantities of viral and cellular RNA were calculated using the standard curve method (112). Viral RNA was normalized to 18S RNA for each sample (Wuhan V1/18S).

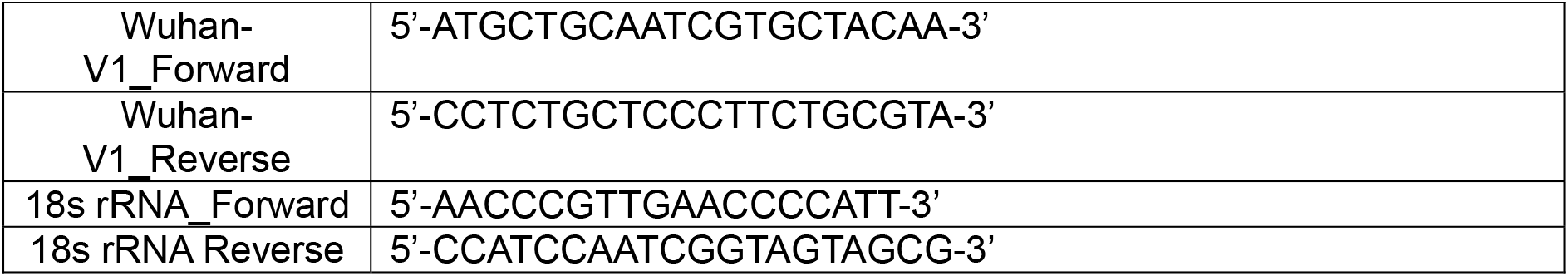

## Acknowledgements

We thank S. Weiss and Y. Li for sharing SARS-Related Coronavirus 2, Isolate USA-WA1/2020 (obtained from the Centers for Disease Control and BEI resources). We thank BEI resources for quantitative SARS-CoV-2 RNA. We thank M. Diamond and S. Hensley for providing anti-Spike antibody (CR3022), C. Coyne for J2 antibody, M. Diamond for oligo sequences. We thank E. Grice for HaCaT cells. We thank C. Kovacsics for Biosafety support. We thank the Cherry lab, the high-throughput screening core, David Roth, and John Epstein for discussions. We thank Timothy Wells and Medicines for Malaria Venture for helpful discussions and compounds. We thank the NIH, Dean’s Innovation Fund, Linda and Laddy Montague, BWF for funding.

## Supplementary Figure Legends

**Figure S1.**
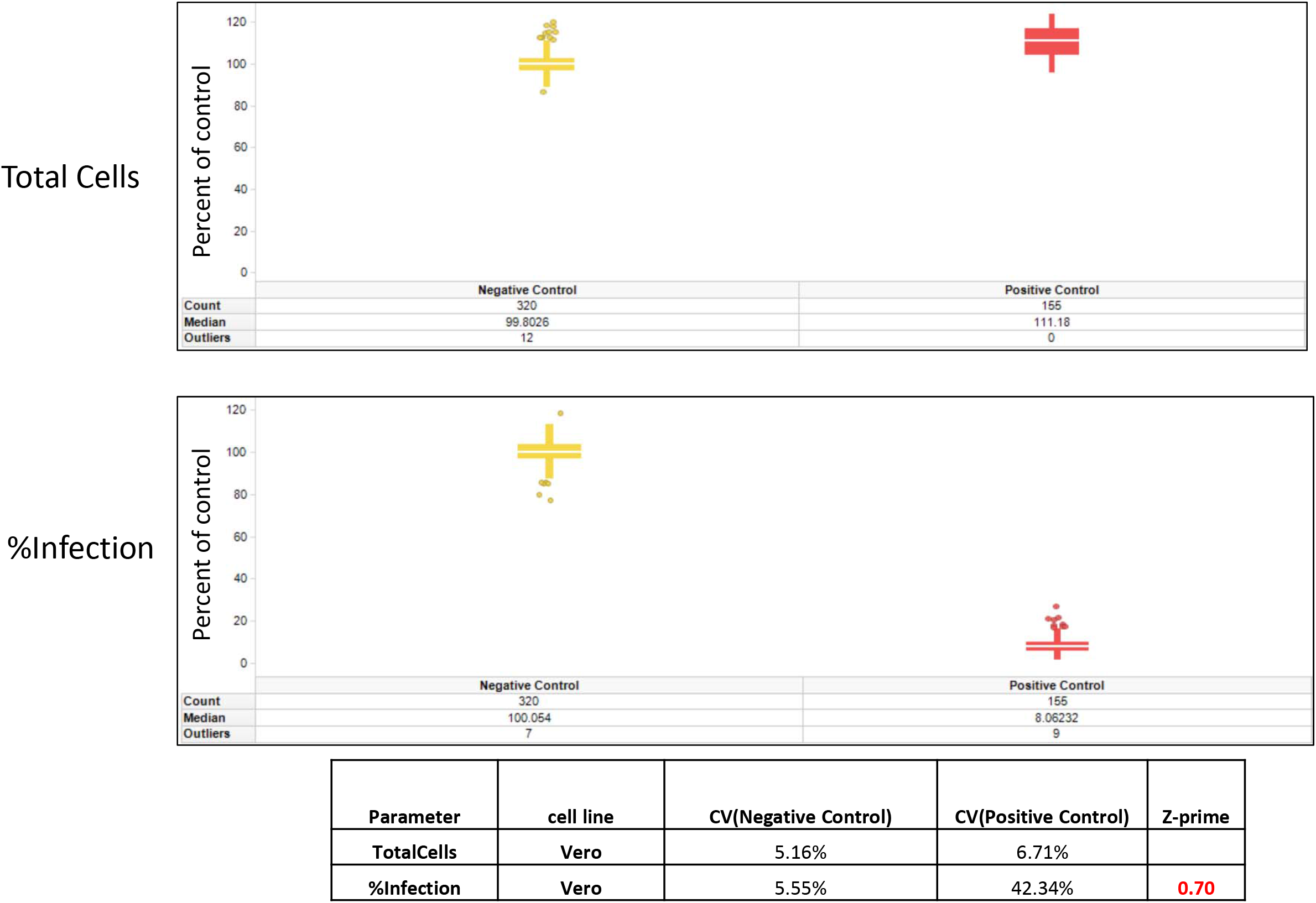
Vero cell screen. Box plots represent the Percentage of Control of Total Cells or Infection after normalization to the average of each plate for all negative (DMSO) or positive (10uM remdesivir) across all screening plates. Horizontal white bar represents the median. Vertical bar represents the standard deviation. Circles represent outlier wells.

**Figure S2.**
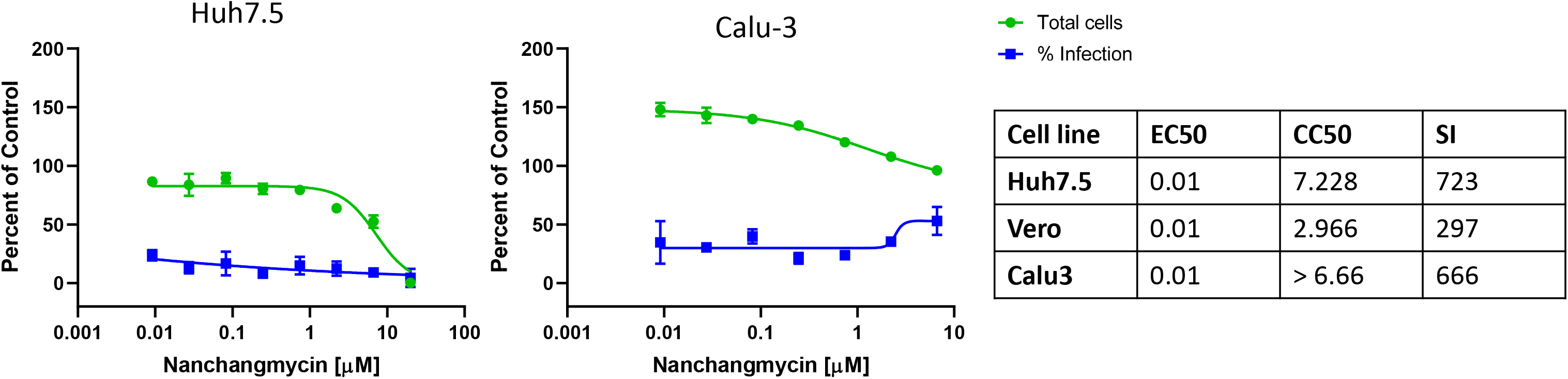
Nanchangmycin is antiviral against SARS-CoV-2 in Huh7.5 cells. Dose-response analysis of nanchangmycin in Huh7.5 cells

**Figure S3.**
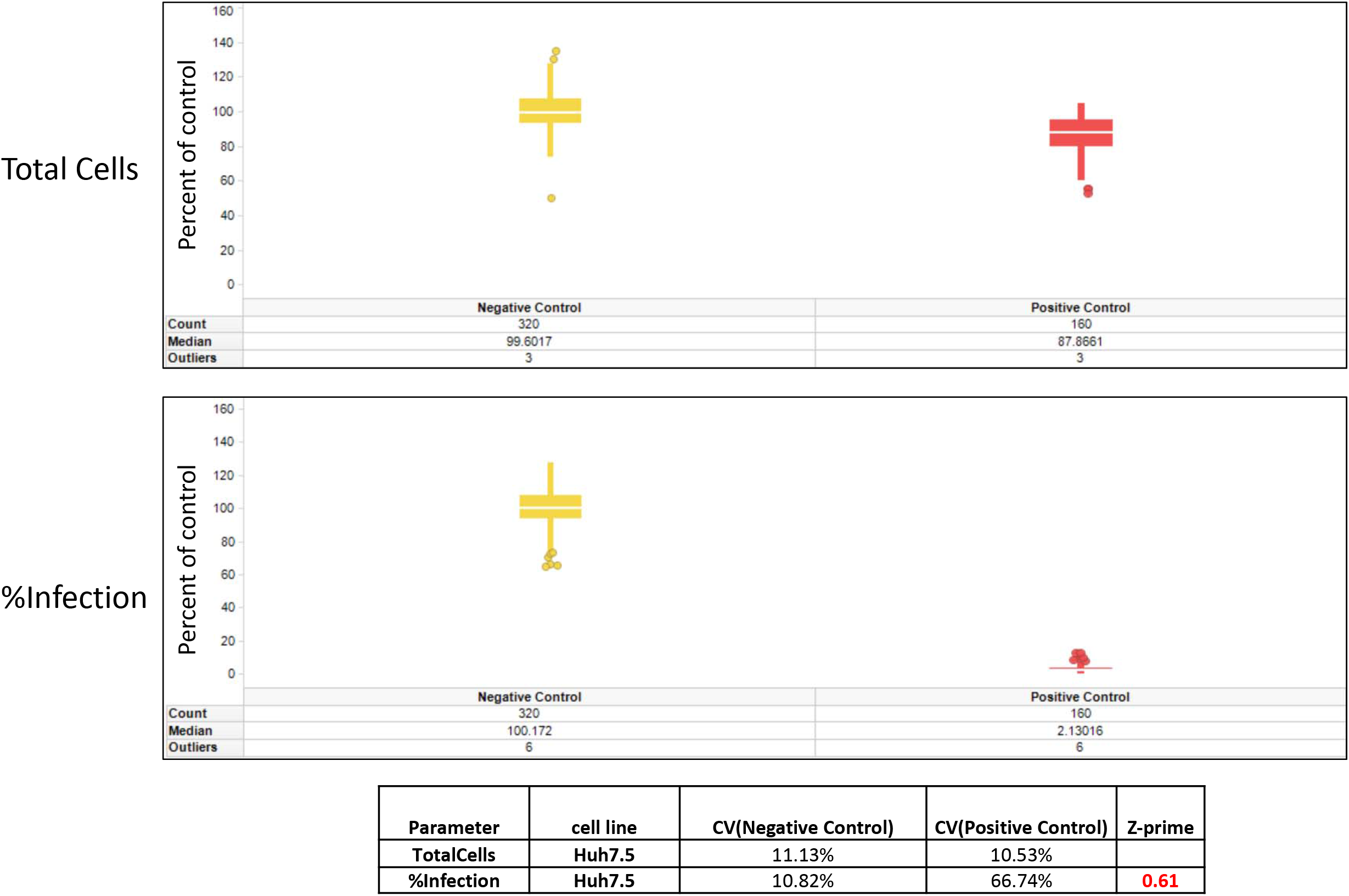
Huh7.5 cell screen. Box plots represent the Percentage of Control of Total Cells or Infection after normalization to the average of each plate for all negative (DMSO) or positive (10uM remdesivir) across all screening plates. Horizontal white bar represents the median. Vertical bar represents the standard deviation. Circles represent outlier wells.

**Figure S4.**
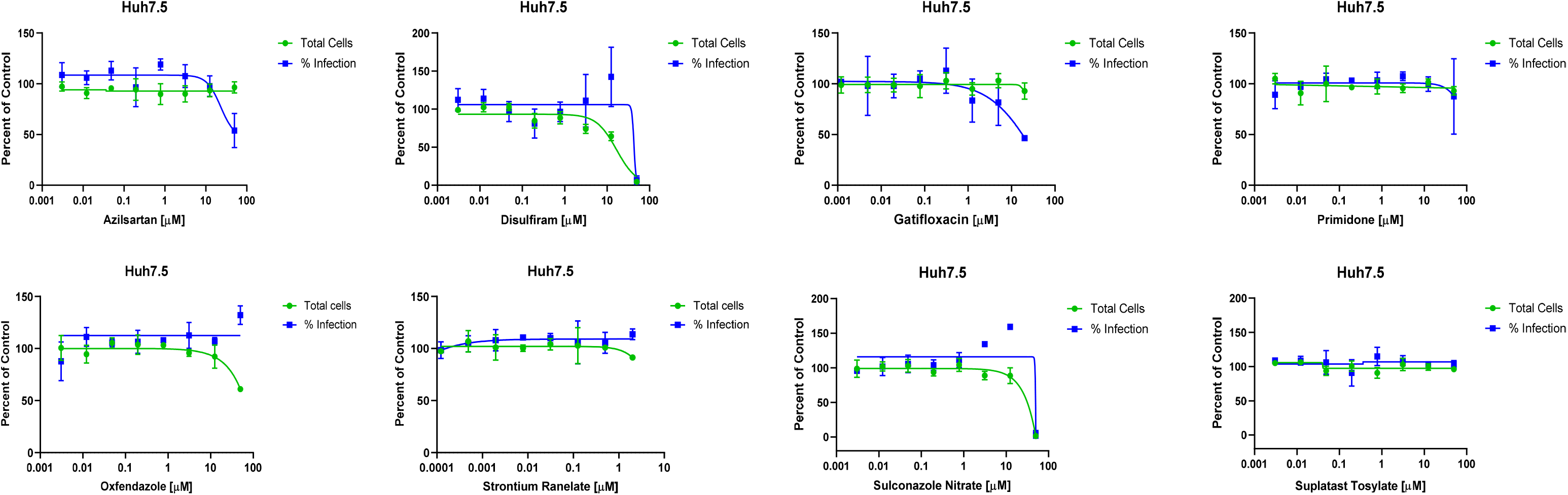
Candidates that did not show antiviral activity in Huh7.5 cells. Dose-response analysis of Huh7.5 cells treated with the indicated drugs.

**Figure S5.**
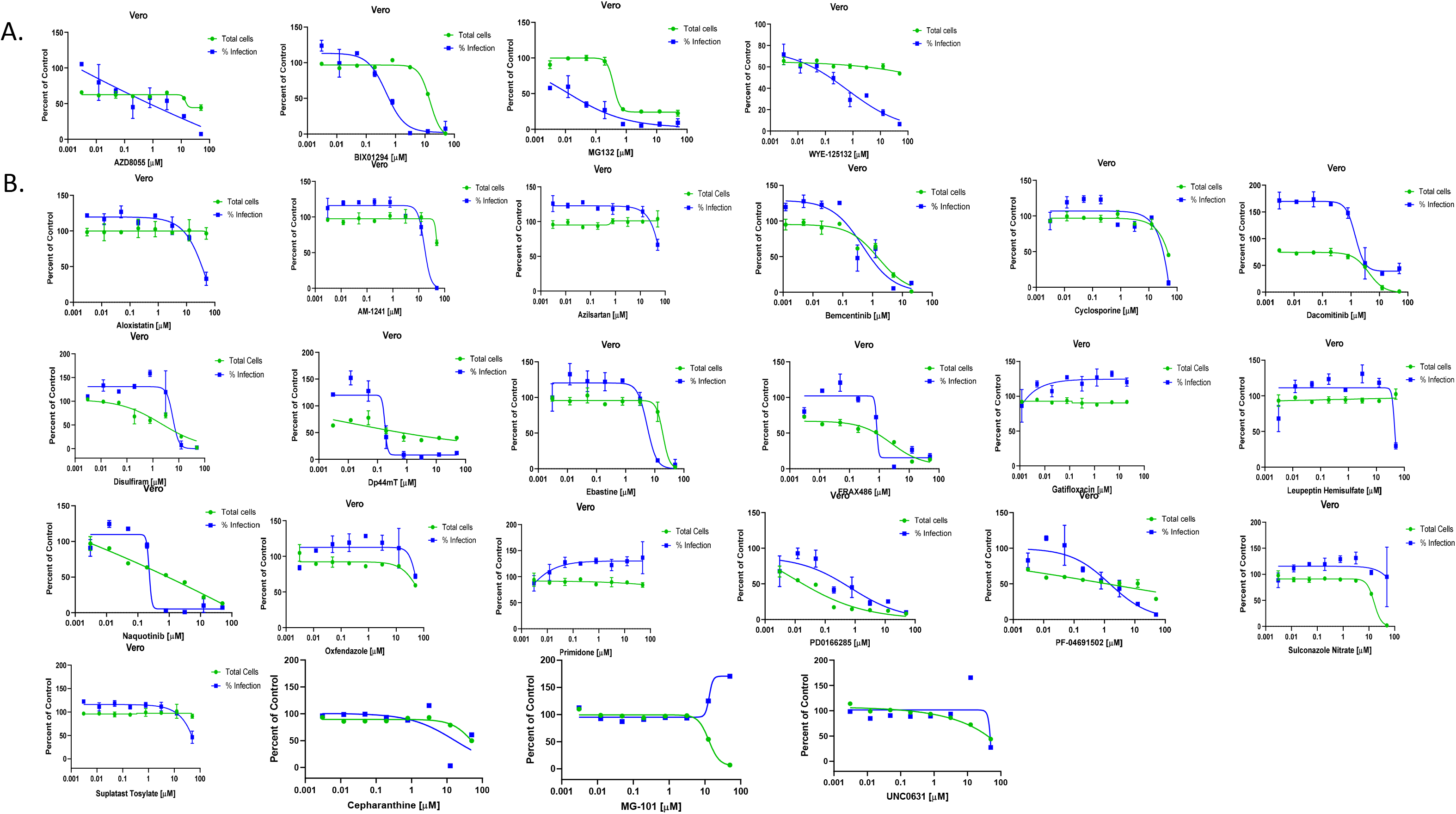
Dose-responses of the candidates identified in Huh7.5 cells in Vero cells. Dose-response analysis Vero cells with the indicated drugs.

**Figure S6.**
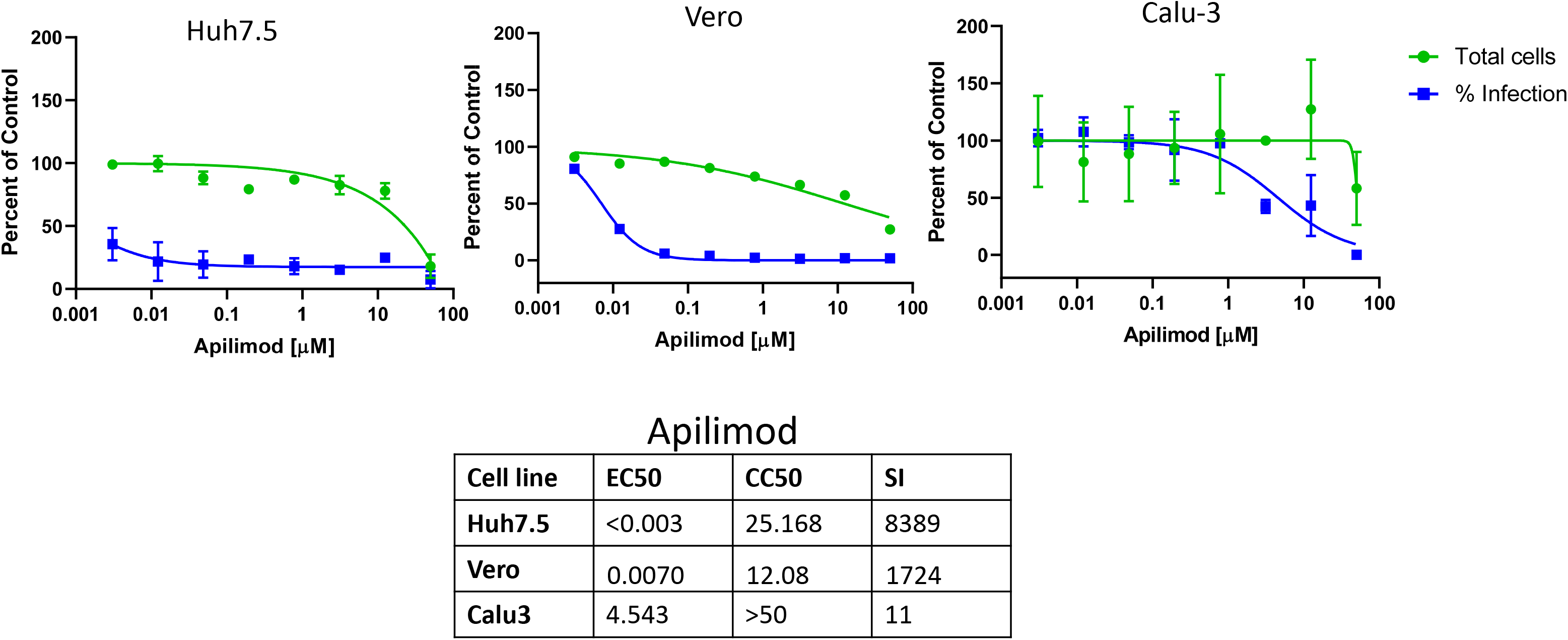
Dose-responses of apilimod across cell types.

**Figure S7.**
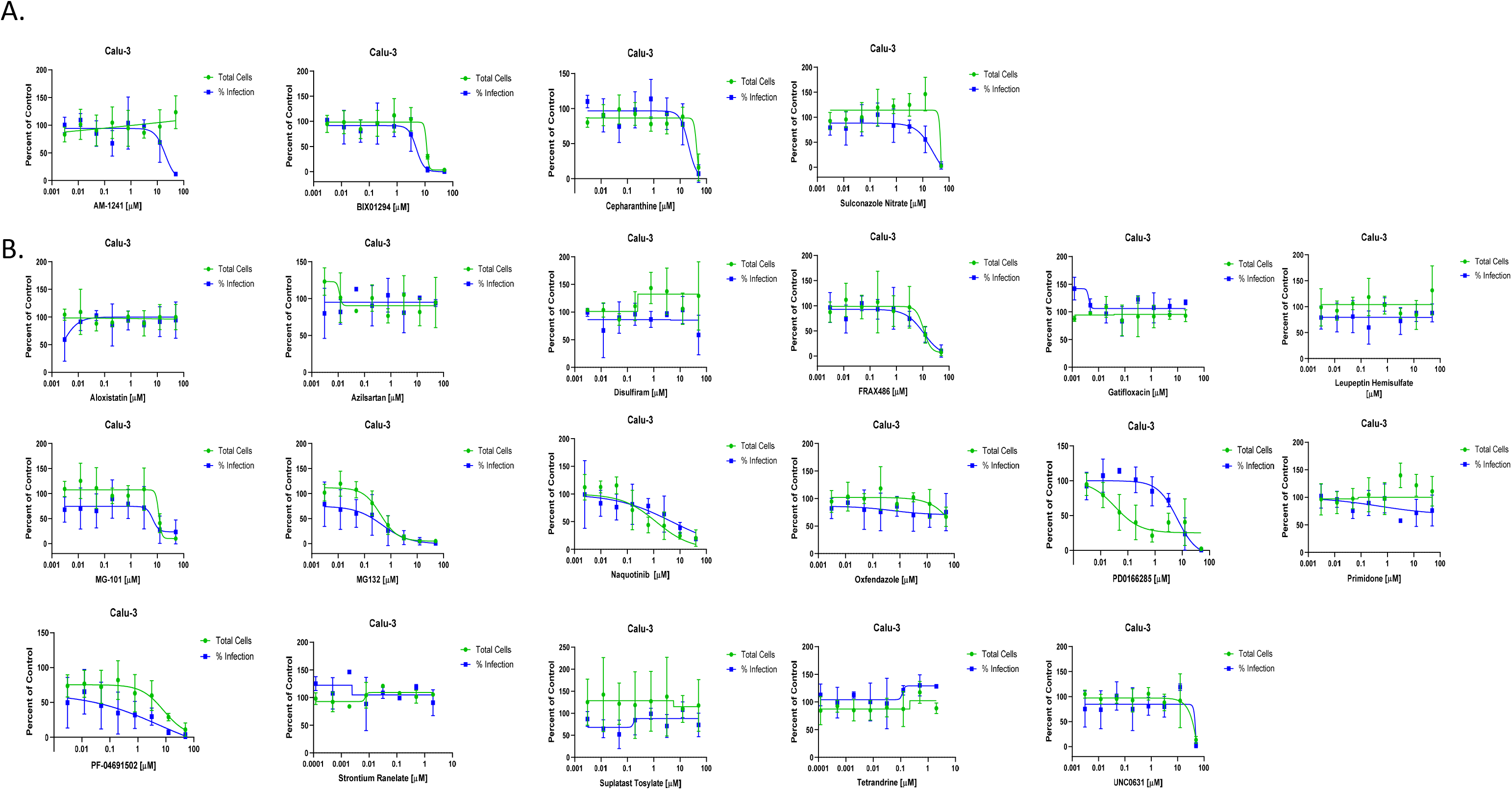
Dose-responses of the candidates identified in Huh7.5 cells in Calu-3 cells. Dose-response analysis of Calu-3 treated with the indicated drugs.

**Figure S8.**
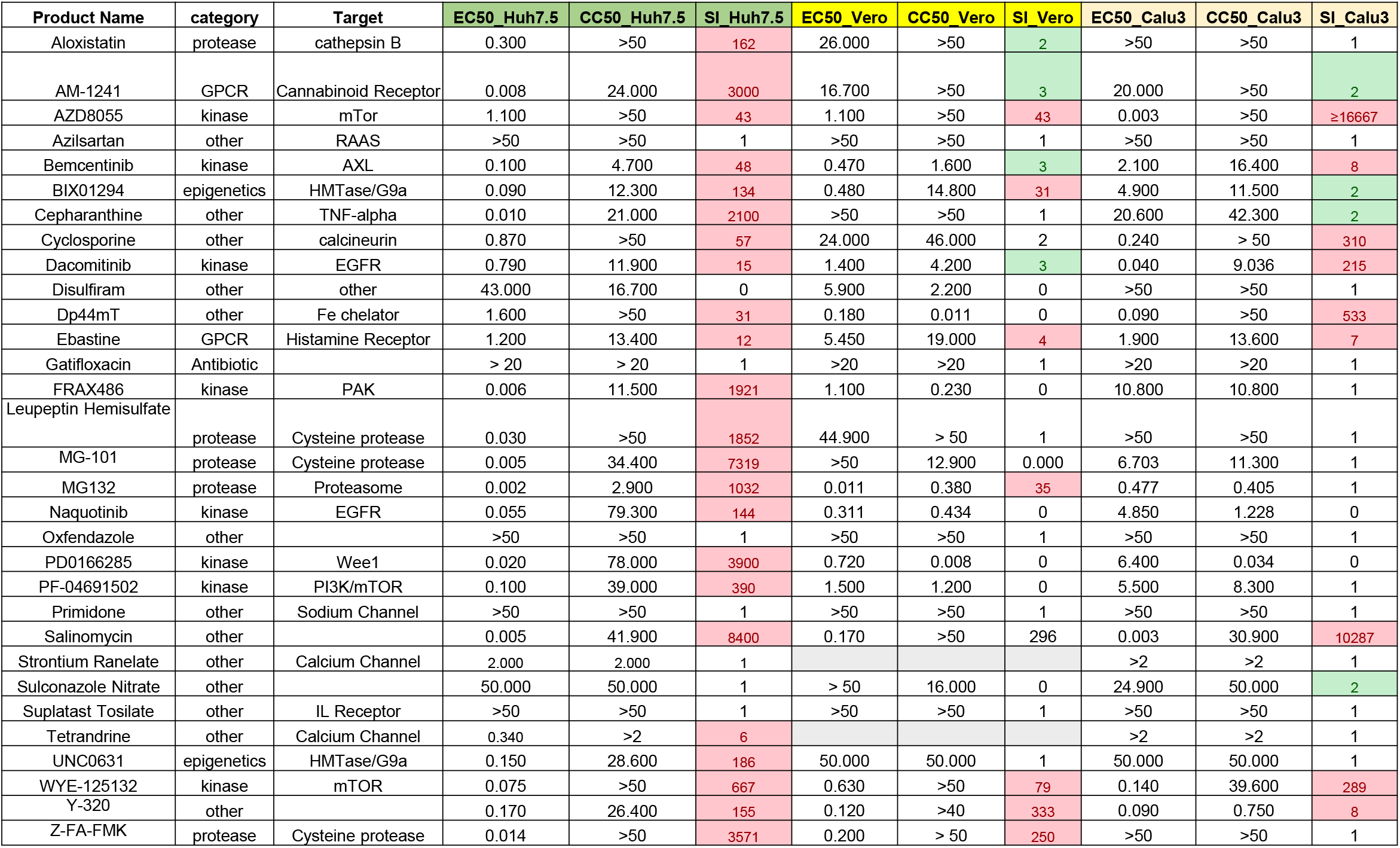
Antivirals identified in Huh7.5 cells. The product name, category and target are shown along with the IC50, CC50 and SI for Huh7.5, Vero and Calu-3 cells.

**Figure S9.**
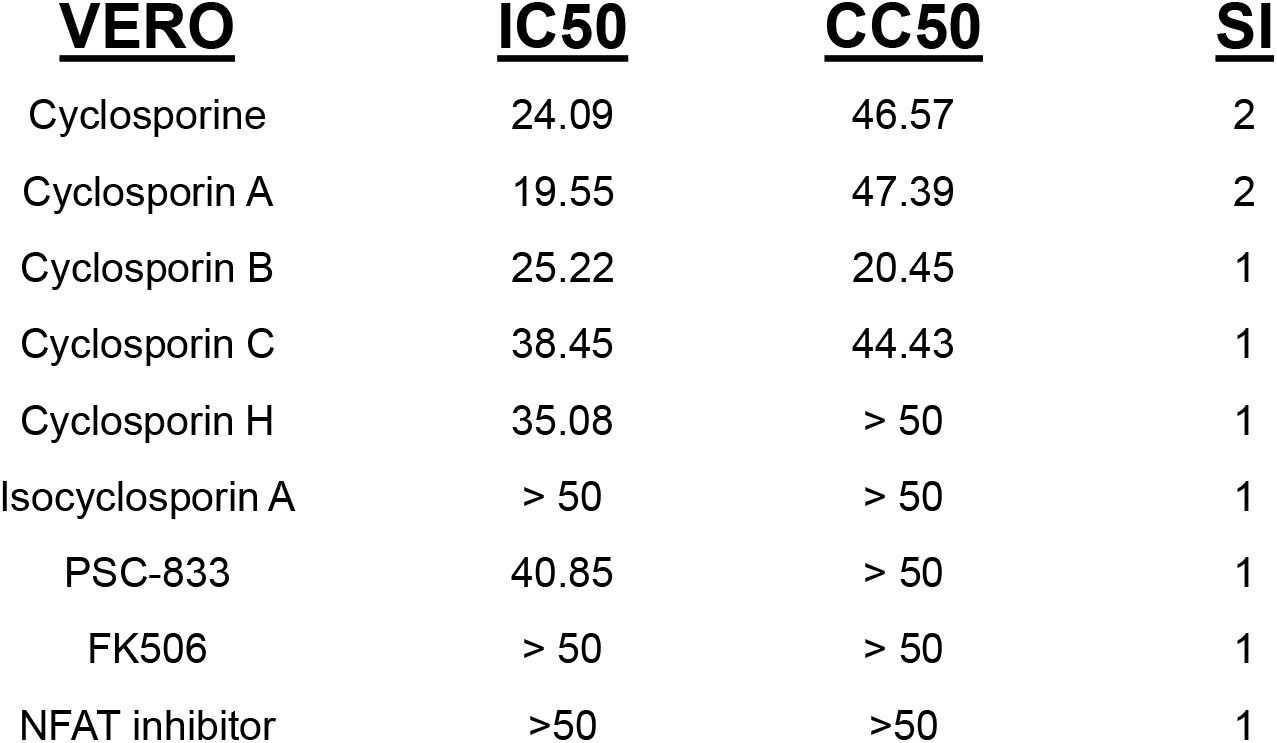
Cyclosporine and drugs with related activities are not antiviral in Vero cells. The drug name along with the IC50, CC50 and SI for Vero cells for the indicated drugs.

---

**Supplementary Table 1**: Primary screen in Vero cells

**Supplementary Table 2**: Primary screen in Huh7.5 cells

**Supplementary Table 3**: COVID-19 Box and antiviral screening data

